# Patch2Self denoising of Diffusion MRI with Self-Supervision and Matrix Sketching

**DOI:** 10.1101/2022.03.15.484539

**Authors:** Shreyas Fadnavis, Agniva Chowdhury, Joshua Batson, Petros Drineas, Eleftherios Garyfallidis

## Abstract

Diffusion-weighted magnetic resonance imaging (DWI) is the only noninvasive method for quantifying microstructure and reconstructing white-matter pathways in the living human brain. Fluctuations from multiple sources create significant additive noise in DWI data which must be suppressed before subsequent microstructure analysis. We introduce a self-supervised learning method for denoising DWI data, Patch2Self (P2S), which uses the entire volume to learn a full-rank locally linear denoiser for that volume. By taking advantage of the oversampled *q*-space of DWI data, P2S can separate structure from noise without requiring an explicit model for either. The setup of P2S however can be resource intensive, both in terms of running time and memory usage, as it uses all voxels (*n*) from all-but-one held-in volumes (*d* − 1) to learn a linear mapping Φ : ℝ^*n*×(*d*−1)^ ↦ ℝ^*n*^ for denoising the held-out volume. We exploit the redundancy imposed by P2S to alleviate its performance issues and inspect regions that influence the noise disproportionately. Specifically we introduce P2S-sketch, which makes a two-fold contribution: *(1)* P2S-sketch uses matrix sketching to perform self-supervised denoising. By solving a sub-problem on a smaller sub-space, so called, *coreset*, we show how P2S can yield a significant speedup in training time while using less memory. *(2)* We show how the so-called statistical leverage scores can be used to *interpret* the denoising of dMRI data, a process that was traditionally treated as a black-box. Our experiments conducted on simulated and real data clearly demonstrate that P2S via matrix sketching (P2S-sketch) does not lead to any loss in denoising quality, while yielding significant speedup and improved memory usage by training on a smaller fraction of the data. With thorough comparisons on real and simulated data, we show that Patch2Self outperforms the current state-of-the-art methods for DWI denoising both in terms of visual conspicuity and downstream modeling tasks. We demonstrate the effectiveness of our approach via multiple quantitative metrics such as fiber bundle coherence, *R*^2^ via cross-validation on model fitting, mean absolute error of DTI residuals across a cohort of sixty subjects.

## 1. Introduction

Diffusion MRI (dMRI) is a 4D acquisition method that generates a series of 3D volumes each corresponding to different gradient directions [6, 44]. Each 3D volume provides unique information about the underlying diffusion processes in the brain. This information is used to probe the tissue microstructure in the living brain by modeling the signal per voxel using a variety of biophysical models [64, 65]. This derived information can however be corrupted due to low signal-to-noise ratio (SNR). Multiple sources of noise are apparent in dMRI that reduce SNR. Furthermore, with new acquisition schemes, high-field MR gradients [80, 61] and multi-dimensional diffusion encoding strategies [71, 41] the effect of noise sources is exaggerated and affects image conspicuity.

In the past, denoising dMRI has been tackled using a variety of methods belonging to different classes based on the signal assumptions imposed. The first class of denoising methods used for DWI data were extensions of techniques developed for 2D images, such as non-local means (NL-means) [17] and its variants [16, 12]), total variation norm minimization [46], cosine transform filtering [57], empirical Bayes [3] and correlation based joint filtering [79]. Some other methods take more direct advantage of the fact that DWI measurements have a special 4D structure, representing many acquisitions of the same 3D volume at different b-values and in different gradient directions. Assuming that small spatial structures are more-or-less consistent across these measurements, these methods project to a local low-rank approximation of the data [66, 58]. The top performing methods are overcomplete Local-PCA (LPCA) [58] and its Marchenko-Pastur extension [81]. The current state-of-the-art unsupervised method for denoising DWI is the Marchenko-Pastur PCA, which handles the choice of rank in a principled way by thresholding based on the eigenspectrum of the expected noise covariance matrix. Note that Marchenko-Pastur PCA, like the classical total variation norm and NL-means methods, requires a noise model (via the Marchencko-Pastur distribution) to do the denoising, either as an explicit standard deviation and covariance as in LPCA, or implicitly in the choice of a noise correction method [47, 76].

**Self-supervised learning**, as a sub-domain of unsupervised learning algorithms, has been rapidly gaining traction over the past years. Novel strategies of self-supervision are being developed and employed for different problems such as multimodal learning [86, 62], self-labeling [56, 48], learning semantic context [19] and contrastive predictive coding [4]. Denoising strategies based on self-supervision have revolutionized image denoising performance across different scientific domains[45, 40]. Leveraging statistical independence of noise, first introduced in the work of Noise2Noise (N2N) [53] was given a theoretical grounding in the work of Noise2Self by posing it under a self-supervised framework [7]. Other approaches similar to N2S and N2N (such as Noise2Void, etc.) were also proposed around the same time [49, 50]. In this work, we introduce **Patch2Self (P2S)** [29] which belongs to this same family of algorithms and leverages the statistical independence of noise. P2S combines N2N and N2S and recasts the 4D self-supervised denoising of dMRI as an image in-painting problem [9]. Instead of holding out a part of the same 3D volume, P2S holds out the entire 3D volume itself and learns to predict a de-noised version of the held-out volume as a linear combination of the remaining 3D volumes. P2S relies on the theory of 𝒥-invariance [7, 29] to perform the denoising, which can be seen as a way of performing 4D image in-painting in the *q*-space [5]. In the case of dMRI data, denoising is typically done on a single subject and using P2S we show how one can get state-of-the-art denoising performance using linear regression as its backbone [13, 31, 75].

Self-supervised denoisers (typically making assumptions on the noise) outperform traditional methods of denoising, but can be computationally expensive as they do not place assumptions on signal properties such as sparsity, compressibility, repetition, etc. In pursuit of acquiring data at a higher resolution and to extract detailed diffusion information, the dimensionality of the data increases rapidly (per-volume and per-scan). Moreover, due to the advent of high-field scanners, more noise is induced in the signal due to the use of stronger magnetic gradients. Along with the dimensionality of a single scan, the number of scans acquired and released for analyses is also increasing rapidly, requiring fast and efficient denoising algorithms for clinical use.

To alleviate these issues, we also introduce **P2S-sketch**, which proposes the sketching of the large matrix 𝒜 [29] constructed for training the denoiser via P2S to create a coreset. 𝒜 is constructed by vectorizing each 3D volume and concatenating it along the columns of 𝒜. This is called a *Casorati matrix* where each independent measurement forms a column of 𝒜 [61]. So, instead of performing the self-supervised denoising on the over-determined set of constraints (𝒜 ∈ ℝ^*n*×*d*^, where *n* ≫ *d*), P2S-sketch samples and rescales the matrix to a much smaller subset of the constraints (𝒜 ∈ ℝ^*s*×*d*^, where *n* ≫ *s* ≈ *d*). By training the self-supervised denoiser on this much smaller induced sub-problem, we show that one can achieve the same level of denoising performance as P2S with a highly reduced time complexity and a much smaller memory footprint. We show the speedup gains both via the theoretical complexity analysis and the empirical comparisons on simulated and real datasets of different dimensionalities. To ensure that P2S-sketch does not hamper the denoising performance, we compare P2S-sketch against P2S on both simulated and real data using the root mean squared error (RMSE) and the *R*^2^ metrics. We also evaluate the performance on the downstream tasks of microstructure modeling and tractography. While the sketch size required for sketching and solving the linear system within P2S-sketch may vary, our experiments suggest at least a 60% redundancy in the training set. We compared the performance of P2S-sketch via different sketching methods such as CountSketch, leverage score sampling, and the Subsampled Randomized Fourier Transform (SRFT). Our results show that leverage score sampling yielded the best performance. We discuss how leverage scores can be used for interpretability of P2S-sketch, revealing which regions of the data have a higher influence on the denoising algorithm. This enables interpretability (crucial to medical imaging) of the self-supervised denoising, which is otherwise treated as a black-box approach. With the help of the Rank Revealing QR (RRQR) decomposition, we also show how one can calibrate the optimal sketch size to construct the coreset for P2S-sketch via a self-supervised loss.

## 2. Methods

In this paper, we introduce two algorithms P2S and P2S-sketch sequentially since P2S-sketch builds on top of P2S-sketch. The organization of the methods is such that each algorithm is preceded by the necessary preliminaries.

### 2.1. Patch2Self: Self-supervised Denoising of Diffusion MRI

#### 2.1.1. Patches and Local Matrix Approximations

Patch-based self-supervision has been used to learn representations that are invariant to distortions [21, 20], for learning relations between patches [19], for filling in missing data (i.e. image in-painting) [67], etc. P2S abides by a similar patch-based approach where we learn an underlying clean signal representation that is invariant to random fluctuations in the observed signal. Inspired by the local matrix approximation works presented in [52, 11], we formulate a global estimator per 3D volume of the 4D data by training on local patches sampled from the remaining volumes. This estimator function, thus has access to local and non-local information to learn the mapping between corrupted signal and true signal, similar to dictionary learning [35, 74, 77, 37, 10] and non-local block matching [18]. Due to the self-supervised formulation, P2S can be viewed as a non-parametric method that regresses over patches from all other volumes except from the one that is held-out for denoising. Our experiments demonstrate that a simplistic linear-regression model can be used to denoise noisy matrices using *p*-neighbourhoods and a class of 𝒥-invariant functions.

### 2.2. Denoising via Self-Supervised Local Approximations

#### Extracting 3D patches

In the first phase of P2S, we extract a *p*-neighbourhood for each voxel from the 4D DWI data. To do so, we construct a 3D block of radius *p* around each voxel, resulting in a local *p*-neighbourhood of dimension *p* × *p* × *p*.

##### Algorithm 1 Patch2Self

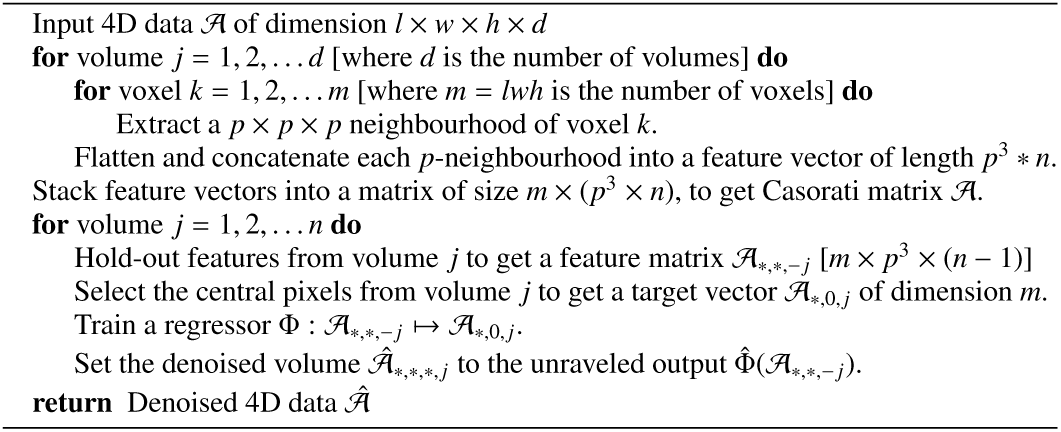

Therefore, if the 4D DWI has *n* volumes {*v*_1_, …, *v*_*n*_} (each volume corresponding to a different gradient direction) and each 3D volume has *m* voxels (see Fig. 1), after extracting the *p*-neighbourhoods, we get a *m* × *p* × *p* × *p* × *n* tensor. Next, we flatten this this tensor along the *p*^*th*^ -dimension to obtain a representation: *m*×(*p*^3^×*n*). Thus, we have transformed the data from the original 4D space to obtain *m* samples of *p*^3^×*n* dimensional 2D feature matrices, which we use for denoising.

**Figure 1:**
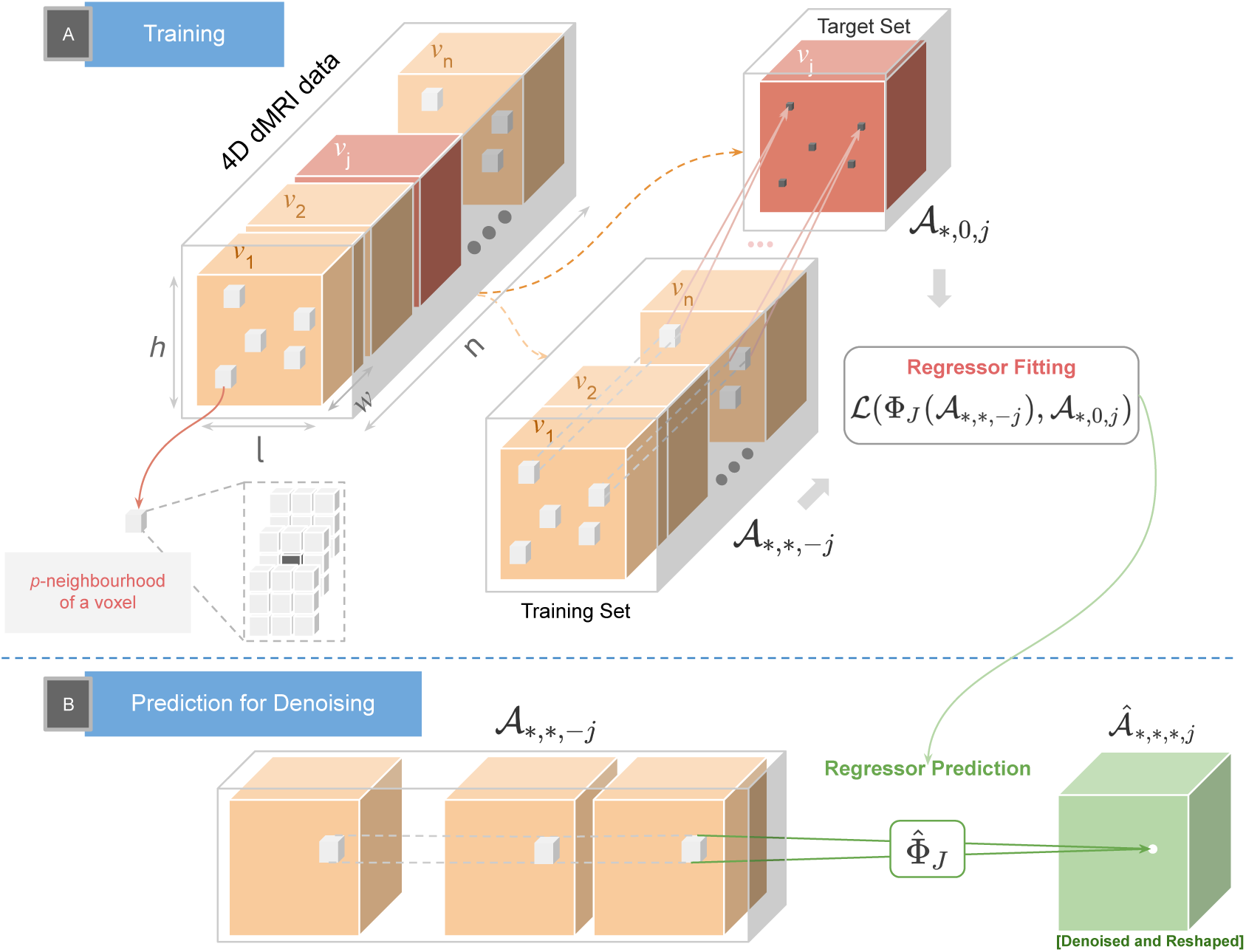
Depicts the workflow of Patch2Self in two phases: **(A)** Is the self-supervised training phase where the 4D DWI data is split into the training 𝒜_∗,∗,− *j*_ and target 𝒜_∗,0, *j*_ sets. *p*-neighbourhoods are extracted from each 3D volume from both 𝒜_∗,∗,− *j*_ and 𝒜_∗,0, *j*_. Φ_*J*_ is the learnt mapping by regressing over *p*-neighbourhoods of 𝒜_∗,∗,− *j*_ to estimate 𝒜_∗,0, *j*_. **(B)** Depicts the voxel-by-voxel denoising phase where 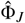 predicts the denoised volume 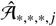 from 𝒜_∗,∗,− *j*_.

#### Self-Supervised Regression

In the second phase, using the *p*-neighbourhoods, P2S reformulates the problem of denoising with a predictive approach. The goal is to iterate and denoise each 3D volume of the noisy 4D DWI data using the following training and prediction phases:

##### (i) Training

To denoise a particular volume, *v*_*j*_, we train the a regression function Φ_*J*_ using *p*-neighbourhoods of the voxels denoted by the set 𝒜. From the first phase, 𝒜 is a set containing *m* training samples with dimension: *p*^3^ × *n*. Next, we hold out the dimension corresponding to volume *v*_*j*_ from each of the *p*-neighbourhoods and use it as a target for training the regressor function Φ_*J*_ (shown in Fig. 1A). Therefore our training set 𝒜_∗,∗,−*j*_ has dimension: *m*×*p*^3^×(*n*−1), where *j* indexes the held out dimension of the *p*-neighbourhoods set. Using the regressor function Φ_*J*_, we use the training set 𝒜_∗,∗,−*j*_ to only predict the center voxel of the set of *p*-neighbourhoods in the corresponding target set of dimension 𝒜_∗,0,−*j*_. The target set, is therefore only an *m*-dimensional vector of the center voxels of the corresponding *p*-neighbourhoods of volume *v*_*j*_. In summary, we use the localized spatial neighbourhood information around each voxel of the set of volumes *v*_−*j*_, to train Φ_*J*_ for approximating the center voxel of the target volume *v*_*j*_. To do so, we propose minimizing the self-supervised loss over the described *p*-neighbourhood sets as follows:

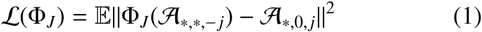

Where, 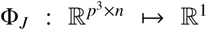, is trained on *m* samples of *p*-neighbourhoods.

##### (ii) Predict

After training for *m* samples, we have now constructed a 𝒥-invariant regressor 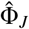 that can be used to de-noise the held out volume *v*_*j*_. To do so, *p*-neighbourhoods from the set 𝒜_∗,∗,−*j*_ are simply fed into 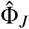 to obtain the de-noised *p*-neighbourhoods corresponding to the denoised volume 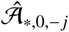. After doing so, for each *j* ∈ {1 … *n*}, we unravel the *p*-neighbourhoods for each volume *v*_*j*_ ∈ {*v*_1_ … *v*_*n*_} (in Fig. 1 as 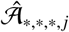) and append them to get denoised 4D DWI data 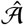 (Details in 1).

###### 𝒥-Invariance

The reason one might expect the regressors learned using the self-supervised loss above to be effective denoisers is the theory of 𝒥-invariance introduced in [7]. Consider the partition of the data into volumes, 𝒥 = {*v*_1_, …, *v*_*n*_}. If the noise in each volume is independent from the noise in each other volume, and a denoising function Φ satisfies the property that the output of Φ in volume *v*_*j*_ does not depend on the input to Φ in volume *v*_*j*_, then according to Proposition 1 of [7], the sum over all volumes of the self-supervised losses in equation 1 will in expectation be equal to the ground-truth loss of the denoiser Φ, plus an additive constant. This means that 𝒥-invariant functions minimizing the self-supervised loss will also minimize the ground-truth loss. This holds by construction for our denoiser Φ = (Φ_1_, …, Φ_*n*_). Intuitively, each Φ_*J*_ only has access to the signal present in the volumes other than *v*_*j*_, and since the noise in those volumes is irrelevant for predicting the noise in *v*_*j*_, it will learn to suppress the fluctuations due to noise while preserving the signal. Note that, if linear regression is used to fit each Φ_*J*_, then the final denoiser Φ is a linear function. Unlike methods which work by thresholding the singular values obtained from a local eigen-decomposition [58, 81], which produce denoised data that are locally low-rank, this mapping Φ can be full-rank.

#### Choice of Regressor

Any class of regressor can be fit in the above method, from simple linear regression/ordinary least squares to regularized linear models like Ridge and Lasso, to more complex nonlinear models. Our code-base allows for the use of any regression model from [68]. Surprisingly, we found that linear regression performed comparably to the more sophisticated models, and was of course faster to train (see supplement for comparisons).

#### Choice of Patch Radius

To determine the effect of changing the patch radius on denoising accuracy, we compute the Root Mean Squared Error (RMSE) between the ground truth and P2S denoised estimates at SNR 15 (details of simulation in supplement). For patch radius zero and one, we show the effect at different number of volumes. The line-plot trend in supplement shows that the difference in the RMSE scores between the two patch radii steadily decreases with an increase in number of volumes. However, with lesser number of volumes, a bigger patch-radius must be used. In the remainder of the text, we use and show results with patch radius zero and linear regressors.

To summarize, leveraging the fact that each 3D volume of the 4D data can be assumed to be an independent measurement of the same underlying object, P2S proposes constructing a large Casorati matrix wherein each column can be assumed to be linearly independent of the other columns. P2S sets up the self-supervised regression task so that each column is denoised by representing it as a combination of the remaining columns. For the sake of simplicity, we denote the self-supervised loss of P2S in the following form for the remainder of the text: 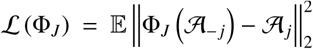. Here 𝒜_*j*_ refers to the voxels corresponding to the volume that were held-out and used as target for training the rest of the voxels from the remaining 3D volumes 𝒜_−*j*_. P2S showed that this 𝒥-invariant function was in fact a linear map Φ_*J*_ : 𝒜_−*j*_ ↦ *𝒜*_*j*_ that achieved the optimal denoising performance. This allows re-writing the P2S as a linear regression problem:

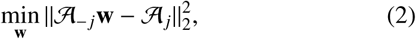

where 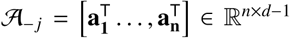 are the features of the design matrix, excluding the held-out *j*-th column. The target for learning Φ_*J*_ is the held out 3D volume 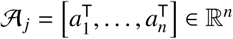.

### 2.3. Coresets for Regression via Matrix Sketching

The self-supervised denoising performed in P2S relies on least squares regression as described in Sec. Preliminaries and Approach. Linear regression typically performed via Cholesky, SVD or QR decomposition needs 𝒪 (*nd*^2^) time. In the case of P2S, the problem setup via the Casorati matrix 𝒜 is massively over-constrained, i.e., *n* ≫ *d*, and has an added time complexity, since the regression is performed *d* times given that each volume needs to be separately denoised. Therefore, in P2S-sketch, instead of computing the exact solution vector 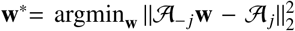, we propose to approximate it using tools from randomized matrix multiplication and sub-space embeddings. The key idea here is to solve a sub-problem 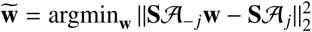 such that

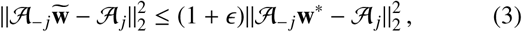

where ϵ is the desired level of accuracy and **S** ∈ ℝ^*s*×*n*^ with *d* ≈ *s* ≪ *n* is the so-called *sketching matrix*. Given the linear sketch **S** 𝒜_−*j*_ of 𝒜_−*j*_, note that computing 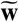 takes 𝒪(*sd*^2^) time, which is indeed much faster than the classical computation of **w**^∗^. Therefore, our goal is to work with a suitable **S** such that the sketch **S**𝒜_−*j*_ can be computed efficiently and satisfies eqn. (3) with high probability. There are several such choices for **S**:

#### Count-sketch [14]

In this case, **S** is a sparse embedding matrix with 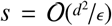 rows and has exactly one non-zero entry per column, which is chosen randomly and set to ±1 independently. The product **S𝒜**_−*j*_ can be computed in time 𝒪(nnz(𝒜_−*j*_)). Assuming the failure probability to be a constant, the overall running time to compute 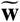 is given by 𝒪(nnz(𝒜_− *j*_))+ poly(*d*/ϵ). Here, nnz(·) denotes the sparsity of the underlying matrix.

#### Fast Johnson-Lindenstrauss transformations [27, 15]

Other choices for the sketching matrix **S** include structured random matrices such as the subsampled randomized Fourier transform (SRFT) or the subsampled randomized Hadamard transform (SRHT). In these cases, the sketching matrix **S** is typically of the form 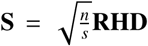, where **D** ∈ ℝ^*n*×*n*^ is a random diagonal matrix with the entries set to ±1 independently; **R** ∈ ℝ^*s*×*n*^ is a subset of *s* rows of the *n* × *n* identity matrix chosen uniformly at random, independently, without replacement; and **H** ∈ ℝ^*n*×*n*^ is either a normalized discrete Fourier transform (for SRFT) or a normalized Walsh-Hadamard matrix (for SRHT). Note that both SRFT and SRHT are based on randomized linear transformations, which can be applied rapidly to arbitrary vectors. Indeed, we can compute the matrix-matrix product **S 𝒜**_−*j*_ in 𝒪(*nd* log *n*) time exploiting the structure of the underlying Fourier/Hadamard matrix; if *s* = 𝒪(*d* + log ^1^/_*ϵ*_ log ^*d*^/_*ϵ*_), then the resulting sketching matrix satisfies eqn. (3). The overall running time is 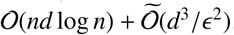.

#### Sampling-based sketching [26]

A third way of achieving a subspace embedding that satisfies eqn. (3) is data-dependent, and can be obtained by sampling rows of a matrix proportional to their leverage scores 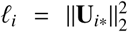 for *i* = 1 … *n*, where **U**_*i*∗_ ∈ ℝ^*n*^ is the *i*-th row of the matrix of the left singular vectors of 𝒜_−*j*_ that are computed using the thin SVD of 𝒜_−*j*_. In this context, the sketching matrix **S** is the so-called *sampling-and-rescaling* matrix of [22] with the sampling probabilities 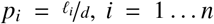, and the sample complexity is given by 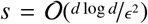. We note that computing *ℓ*_*i*_ s exactly needs access to the matrix **U** which is expensive. Therefore, in practice, approximate leverage scores also suffice and they can be efficiently computed without computing the matrix **U** [23, 14]. We also note that there is another line of work [2, 73] that used sketching as a randomized preconditioner to come up with high precision solutions for overconstrained regression problems. However, in context of P2S-sketch, we can achieve the desired accuracy in denoising performance, even with a *sketch- and-solve* approach as discussed above. Finally, we refer the interested reader to the surveys [85, 25, 54, 24, 60] for background on Randomized Linear Algebra.

## 3. Statistical Leverage for Interpretability

In the previous section, we described how leverage scores can be used to perform importance sampling in order to generate the coreset for P2S-sketch. Here we show how to use the statistical leverage scores of the underlying linear model for interpreting areas of the data that influence the noise. From a statistical perspective, an alternative formulation of leverage scores are the diagonal entries of the projection matrix constructed to solve the linear regression [24, 55]. Typically, one looks at the standard deviation of the noise derived from the models used to perform the denoising [81]. Since P2S and consequently P2S-sketch are set up to denoise in a predictive setting via self-supervision, we can use statistical leverage to get a more detailed view of the factors that would influence the de-noising performance. In Fig. 3A, we show the leverage scores computed on a subject from the PPMI dataset [59]. As one can see in the plot, some voxels in the data exhibit considerably higher statistical leverage. Voxels where an anatomical signal from the brain is captured exhibit higher leverage when compared to the voxels in the background, which have very small leverage scores. In Fig. 3B, we show the spatial map of leverage scores for two example datasets, PPMI and HCP 7T [80]. In the case of the PPMI data, we note that the regions of the white matter, such as the corpus callosum, have higher statistical leverage compared to the rest of the brain. Strikingly, we also see that a slanting structured pattern (indicated with red arrows) appears in the leverage scores map of the HCP 7T data. This structure can also be seen in the noise map of Fig. 4A in the residual map. Leverage score maps of the same HCP 7T subject also highlighted a structured artefact at the bottom of the axial slice (depicted as a white-dotted bounding box). It could be a ghost or a structured artefact added to the 7T data.

**Figure 2:**
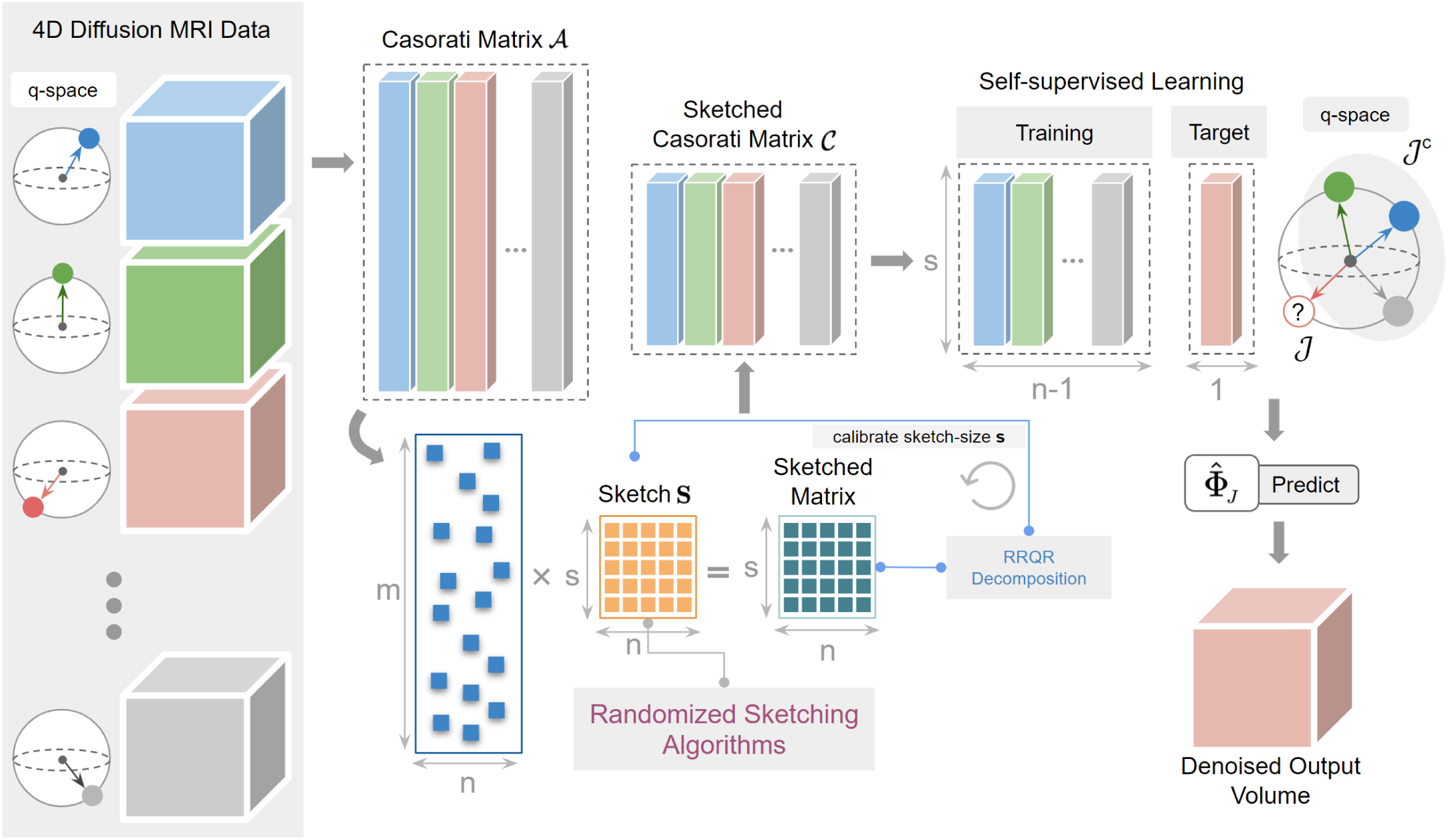
We show how Patch2Self2 works in the case of Diffusion MRI data. The Casorati Matrix 𝒜 is constructed by flattening each gradient direction. 𝒜 is then sketched via a randomized algorithm to generate the smaller sketched Casorati Matrix 𝒞. The 𝒥-invariant training is then performed on 𝒞 to learn 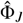 and predict the denoised volume.

**Figure 3:**
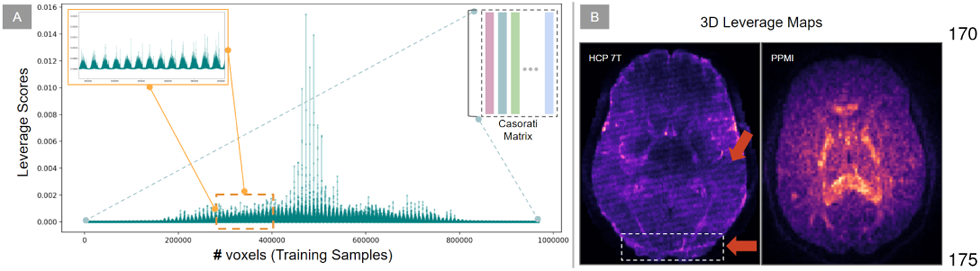
(A) We depict an exemplary point-plot of the row leverage scores computed on the Casorati matrix of a real dataset. (B) We show the axial slice of the 3D leverage score maps computed on two example datasets. Notice that the leverage score maps capture structured artefacts spatially correlated across gradient directions (indicated by red arrows on the HCP 7T data).

**Figure 4:**
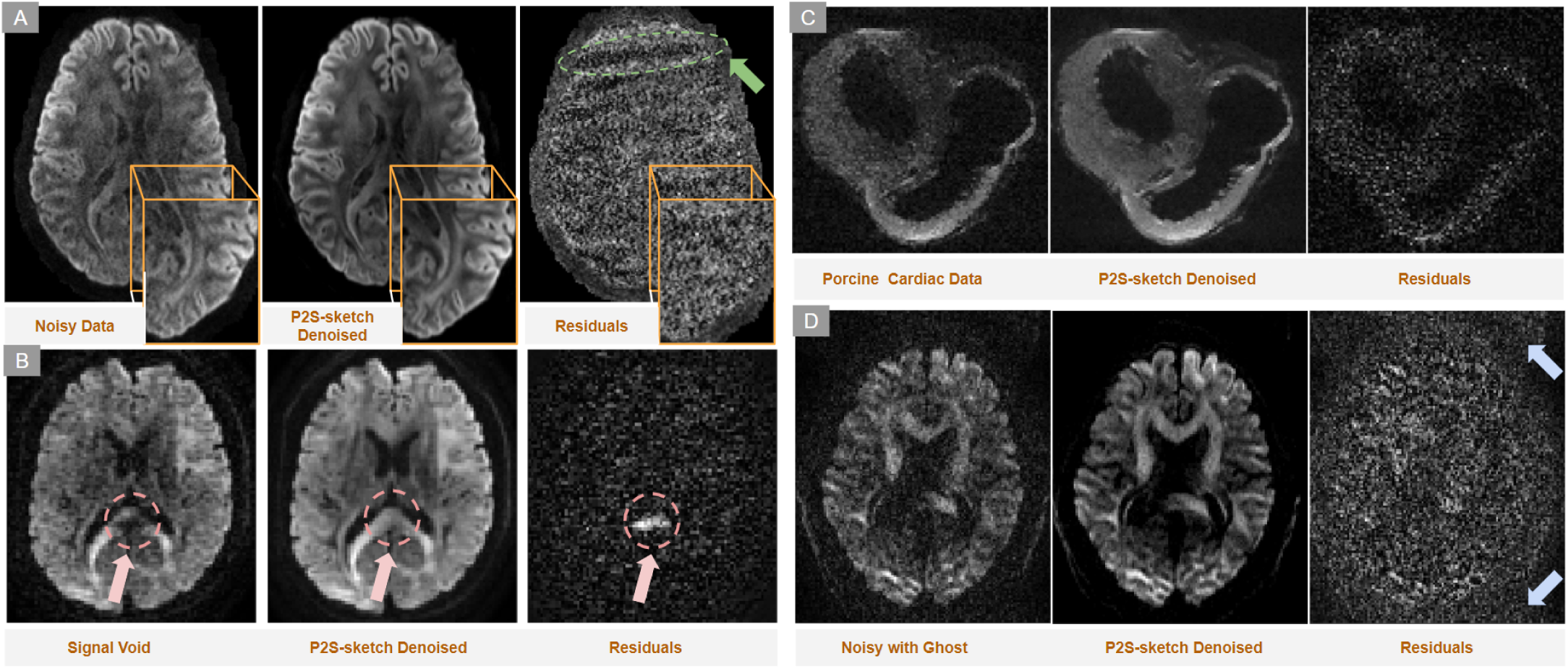
Depicts (A) Noisy band-like pattern in the residual maps suppressed via P2S-sketch (indicated by a green arrow). (B) In-painting of a signal void present in the original data and the predicted signal in the corresponding residual map (indicated by red arrows). (C) Capability of extending P2S-sketch to other organs such as the heart (Porcine Cardiac Data). (D) Suppression of ghosting artefacts (indicated by blue arrows).

## 4. P2S-sketch Algorithm

P2S-sketch extends the idea of P2S (see Sec. Preliminaries and Approach) by performing self-supervised training on coresets. In case of dMRI, we have *d* number of 3D volumes each with dimensionality: *l* × *w* × *h*. Each of the volumes is flattened to a 1D array (*n* = *l* × *w* × *h*) to form a column of the Casorati matrix 𝒜 ∈ ℝ^*n*×*d*^. Next, we sketch this matrix using the sketching matrix **S**(∈ ℝ^*s*×*n*^) to get a sketched Casorati matrix 𝒞 = **S𝒜** (see Sec. Coresets for Regression via Matrix Sketching). We perform a self supervised denoising on this sketched matrix 𝒞 by solving the sub-problem: 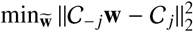. This denoising is performed on a volume-by-volume basis as proposed in P2S, where *j* corresponds to the volume held out for denoising. Thus, P2S-sketch learns a linear map Φ_*J*_ : ℝ^*s*×(*d*−1)^ ↦ ℝ^*s*^, which is a much smaller sub-problem to solve since *s* ≪ *n*. After the training is done, the approximate solution vector 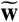 learned via Φ_*J*_ is used to predict the held out volume of 𝒜_*j*_. In order to predict the denoised volume, the full Casorati matrix 𝒜_−*j*_ with all rows is given as input to the function Φ_*J*_ (Algorithm is further detailed in 2). In P2S-sketch, we allow switching between different sketching methods such as SRFT, lever-age scores, and CountSketch (see Sec. Coresets for Regression via Matrix Sketching for details of each). Our results show that sketching via leverage scores outperforms other sketching methods (detailed comparison in Sec. Performance Comparison of Sketching Methods). As per the above procedure, P2S-sketch introduces using a new hyperparameter - the sketch size that needs to be tuned. We propose a 𝒥-invariant self-supervised calibration procedure to find an optimal sketch size *s* based on the QR decomposition and leverage score sketching. Leverage scores are a univariate statistic. When two rows of 𝒜 have identical or similar leverage scores, it implies that the rows are highly correlated and therefore redundant in the construction of the sketched Casorati matrix 𝒞. Thus, a redundancy removal step is often useful to reduce the sketch size *s* while retaining denoising performance. Towards that end, P2S-sketch employs the Rank Revealing QR (RRQR) factorization [38], to flag such redundancies by ranking the rows of the matrix in order of independence. In other words, highly linearly independent rows are given priority after the RRQR has been computed. In P2S-sketch, to calibrate the sketch size, we start with *s* < 50% of the number of rows in 𝒜 and compute the rank-revealing QR (RRQR) decomposition [38] of 𝒞. The RRQR ranks the rows of 𝒞 in order of importance and can be used to remove redundant rows from the sketched matrix. Next, as shown in Fig. 5A, we select the top *k*-ranked rows from 𝒞 and compute the self-supervised loss. Gradually, by increasing the size of *k* in each iteration we compute the self-supervised loss for each iteration. The self-supervised loss for each value of *k* is computed as: ∥Φ_*J*_(𝒜_−*j*_) − 𝒜_*j*_∥. As shown in Fig. 5B, the loss eventually converges to a minimum with minimal change in the denoising performance. At that point, the elbow in the error plot (shown by the red box) reveals the number of redundant rows of the data, which allows us to estimate the (approximately) optimal *s*.

**Figure 5:**
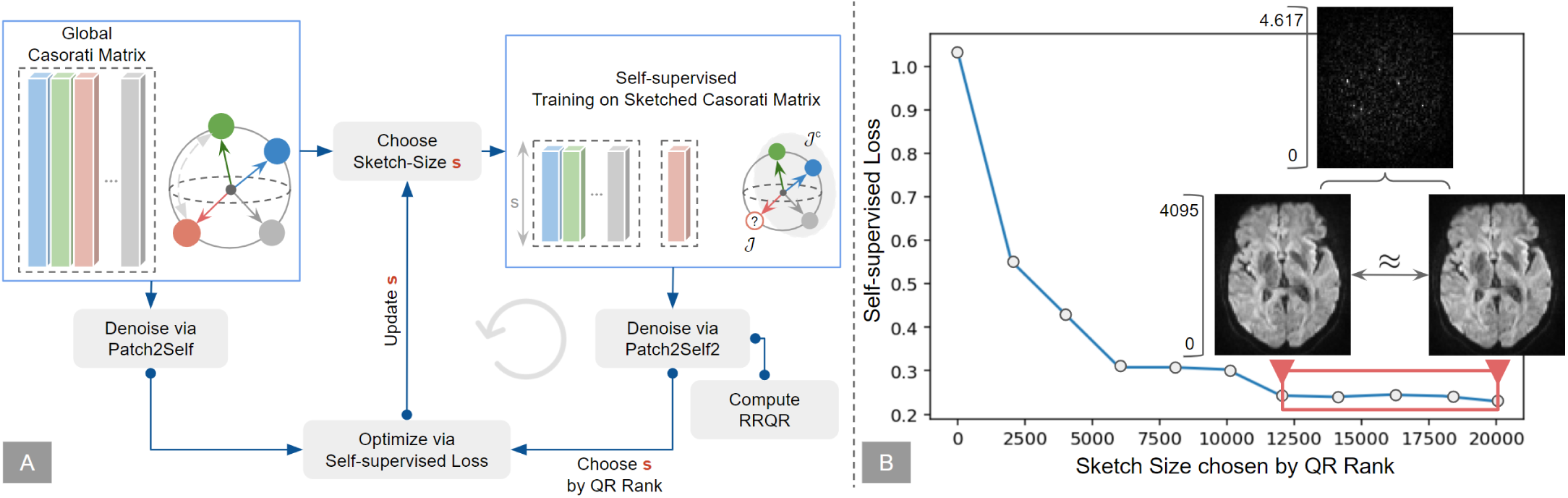
(A) Iterative self-supervised calibration procedure to obtain the optimal sketch size *s* to be used within P2S-sketch using a RRQR decomposition. the self-supervised loss is computed, by iteratively reducing the size of *s* until convergence. (B) We depict an example loss plot of the self-supervised calibration with exemplary PPMI data.

### Leverage Score Sampling Strategies

We can obtain the sketched matrix 𝒞 via leverage score sampling using the following two procedures: (1) Deterministically choosing top *s* lever-age scores; (2) Randomized sampling based on leverage scores. From Fig. 3A, it is evident that only a few rows of the Casorati matrix have a very high leverage score in comparison with the remaining ones. These rows seem to have a higher influence on the denoising performance. As shown in Fig. 6A, the deterministic selection of the highest leverage scores consistently performs worse at all sketch sizes when compared to randomly sampling leverage scores. To investigate this effect, we compared the distribution of the rows corresponding to deterministically chosen top 20K leverage scores against the randomly sampled 20K rows. From the joint plot obtained by fitting a kernel density estimation shown in Fig. 6B, it can be seen that the distribution of the randomly sampled leverage scores forms a bi-modal distribution as opposed to the uni-modal distribution obtained from a deterministic selection of the top 20K leverage scores. This implies that the randomization in the sampling procedure helps denoising by using values that do not have a “high leverage”.

**Figure 6:**
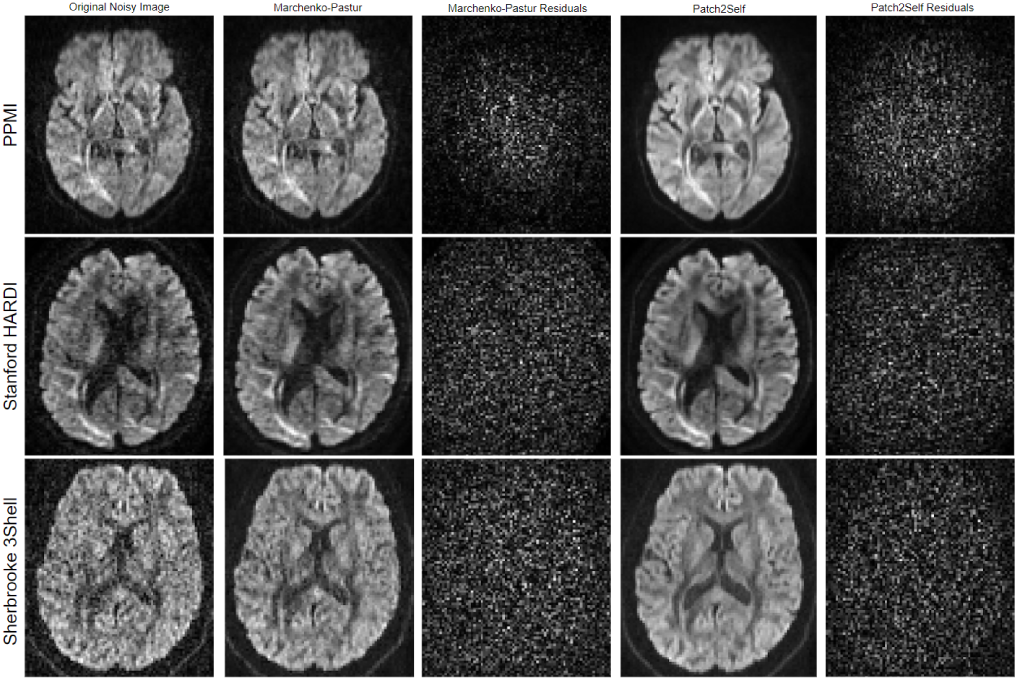
Shows the comparison of denoising on 3 different types of datasets: Parkinson’s Progression Markers Initiative (PPMI), Stanford HARDI and Sher-brooke 3-Shell HARDI data. The denoising of P2S is compared against the original noisy image and Marchenko-Pastur denoised data along with their corresponding residuals. Notice that P2S suppresses more noise and also does not show any anatomical structure in the corresponding residual plots.

## 5. Results

In this section we evaluate P2S and P2S-sketch sequentially since the purpose of evaluation of both algorithms is different. For P2S, we perform qualitative and quantitative comparisons on real and simulated data. While the results on the real data are included in the main text, the results on simulated data for P2S are added to the supplement for compactness and avoiding repetition from [29] and P2S-sketch evaluations. To evaluate the performance on real data, we qualitatively and quantitatively compare the effects of P2S and P2S-sketch denoising on the residual maps, microstructure modeling [65] and tractography [42]. We quantify the effect of different sketch sizes on the speed and accuracy of the approximate solution 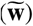 in comparison with the P2S solution (**w**^∗^).

### Algorithm 2 Patch2Self-sketch

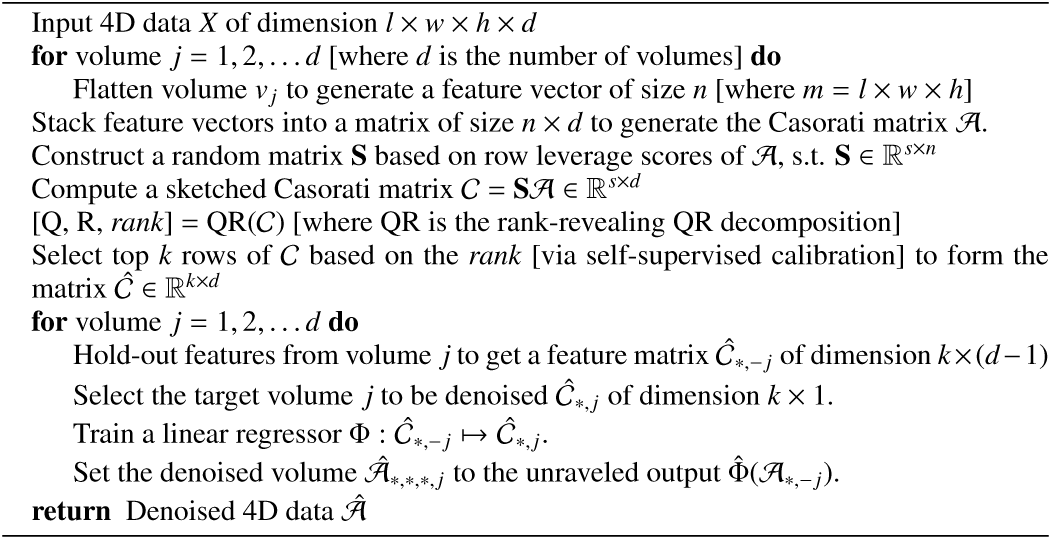

### 5.1. Evaluation of P2S on in-vivo data

We compare the performance of P2S with Marchenko-Pastur on the Parkinson’s Progression Markers Initiative (PPMI) [59], Stanford HARDI [72] and Sherbrooke 3-Shell [33] datasets as shown in Fig. 6. These datasets represent different commonly used acquisition schemes: (1) Single-Shell (PPMI, 65 gradient directions), (2) High-Angular Resolution Diffusion Imaging (Stanford HARDI, 160 gradient directions) and (3) Multi-Shell (Sherbrooke 3-Shell, 193 gradient directions). For each of the datasets, we show the axial slice of a randomly chosen 3D volume and the corresponding residuals (squared differences between the noisy data and the denoised output). Note that both, Marchenko-Pastur and P2S, do not show any anatomical features in the error-residual maps, so it is likely that neither is introducing structural artifacts. P2S produced more visually coherent outputs, which is important as visual inspection is part of clinical diagnosis.

### 5.2. Effect of P2S on Tractography

To reconstruct white-matter pathways in the brain, one integrates orientation information of the underlying axonal bundles (streamlines) obtained by decomposing the signal in each voxel using a microstructure model [8, 65]. Noise that corrupts the acquired DWI may impact the tractography results, leading to spurious streamlines generated by the tracking algorithm. We evaluate the effects of denoising on probabilistic tracking [34] using the Fiber Bundle Coherency (FBC) metric [69]. To perform the probabilistic tracking, the data was first fitted with the Constant Solid Angle (CSA) model [1]. The Generalized Fractional Anisotropy (GFA) metric extracted from this fitting was used as a stopping criterion within the probabilistic tracking algorithm. The fiber orientation distribution information required to perform the tracking was obtained from the Constrained Spherical Deconvolution (CSD) [78] model fitted to the same data. In Fig. 7A, we show the effect of de-noising on tractography for the Optic Radiation (OR) bundle as in [69]. The OR fiber bundle, which connects the visual cortex:V1 (calcarine sulcus) to the lateral geniculate nucleus (LGN), was obtained by selecting a 3 × 3 × 3 Region Of Interest (ROI) using a seeding density of 6. After the streamlines were generated, their coherency was measured with the local FBC algorithm [69, 28]), with *yellow-orange* representing - spurious/incoherent fibers and *red-blue* representing valid/coherent fibers. In Fig, 7, OR bundle tracked from original/ raw data contains 3114 streamlines, Marchenko-Pastur denoised data [81] contains 2331 streamlines and P2S denoised data contains 1622 streamlines. P2S outperforms Marchenko-Pastur by reducing the number of incoherent streamlines, as can be seen in the *red-blue* (depicting high coherence) coloring in Fig. 7A.

**Figure 7:**
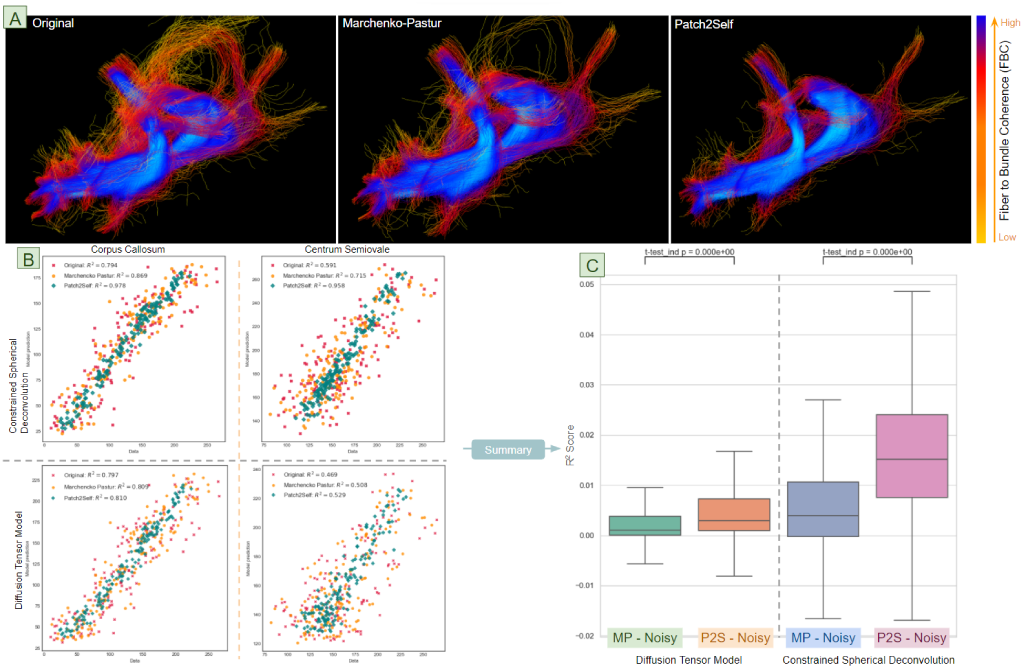
(A) Depicts the Fiber to Bundle Coherency (FBC) density map projected on the streamlines of the optic radiation bundle generated by the probabilistic tracking algorithm. The color of the streamlines depicts the coherency − *yellow* corresponding to incoherent and *blue* corresponding to coherent. Notice that the number of incoherent streamlines present in the original fiber-bundle is reduced after Marchenko-Pastur denoising. P2S denoising further reduces spurious tracts, resulting in a cleaner representation of the fiber bundle. (B) Quantitative comparison of the goodness-of-fit evaluated using a cross-validation approach. Depict the scatter plots of the model predictions obtained by fitting CSD to voxels in the corpus callosum (CC) and centrum semiovale (CSO) for original (noisy), Marchenko-Pastur (denoised) and P2S (denoised) data. Similarly, also show the scatter plots of predictions obtained from DTI fitting in the same voxel locations. Top-left of each plot shows the *R*^2^ metric computed from each model fit on the corresponding data. (C) Box-plots quantifying the increase in *R*^2^ metric after fitting downstream DTI and CSD models. The *R*^2^ improvements in each case are plotted by subtracting the scores of model fitting on noisy data from *R*^2^ of fitting each denoised output. Note that the consistency of microstructure model fitting on P2S denoised data is higher than that obtained from Marchenko-Pastur (see 5.3 for details and significance).

### 5.3. Impact of P2S on Microstructure Model Fitting

The domain of microstructure modeling employs either mechanistic or phenomenological approaches to resolve tissue structure at a sub-voxel scale. Fitting these models to the data is a hard inverse problem and often leads to degenerate parameter estimates due to the low SNR of DWI acquisitions [64]. We apply two of the most commonly used diffusion microstructure models, Constrained Spherical Deconvolution (CSD) [78] and Diffusion Tensor Imaging (DTI) [6], on raw and denosied data. DTI is a simpler model that captures the local diffusion information within each voxel by modeling it in the form of a 6-parameter tensor. CSD is a more complex model using a spherical harmonic representation of the underlying fiber orientation distributions. In order to compare the goodness of each fit, we perform a k-fold cross-validation (CV) [39] at two exemplary voxel locations, corpus callosum (CC), a single-fiber structure, and centrum semiovale (CSO), a crossing-fiber structure. The data is divided into *k* = 3 different subsets for the selected voxels, and data from two folds are used to fit the model, which predicts the data on the held-out fold. The scatter plots of CV predictions against the original data are shown in Fig. 7B for those two voxels. As measured by *R*^2^, P2S has a better goodness-of-fit than Marchenko-Pastur by 22% for CC and 65% for CSO. To show that P2S consistently improves model fitting across all voxels, in Fig. 7C we depict the improvement of the *R*^2^ metric obtained from the same procedure for the axial slice (4606 voxels) of masked data (using [72] data). This was done by simply subtracting the goodness-of-fit *R*^2^ scores of fitting noisy data, from Marchenko-Pastur and P2S denoised data for both CSD and DTI models. P2S shows a significant improvement on both DTI and CSD (two-sided t-test, *p* < 1e-300, Fig. 7C). In order to evaluate the improvement of model fitting, we compared the residuals after fitting the DTI model both before and after denoising. To do so, we made use of 60 subjects from the LA5c cohort (30 controls and 30 schizophrenics) and compared the improvement of residuals after denoising and fitting the DTI model. As one can see via Fig. 10, P2S reduces the residual error much more than MPPCA. The comparison was done by computing the mean absolute error of the residuals between the raw and denoised images to quantify the improvement of model fitting after denoising. Now that we have shown of the noise suppression of P2S, we shift the focus towards P2S-sketch where we start with evaluating different sketching methods and benchmark its performance against P2S. We adopt the same evaluation scheme as P2S [30] where we show similar performance of P2S-sketch against P2S-sketch on denoising performance, microstructure and tractography.

**Table. 1:** Quantitative comparison of Raw, P2S and P2S-sketch denoised data via MSE and *R*^2^ metrics.

| Data | SNR 5 |  | SNR 10 |  | SNR 15 |  | SNR 20 |  | SNR 25 |  |
| --- | --- | --- | --- | --- | --- | --- | --- | --- | --- | --- |
| | MSE | $R^2$ | MSE | $R^2$ | MSE | $R^2$ | MSE | $R^2$ | MSE | $R^2$ |
| Raw | 4.45 | 0.01 | 2.39 | 0.15 | 1.65 | 0.37 | 1.32 | 0.56 | 1.16 | 0.69 |
| P2S | 2.98 | 0.17 | 1.43 | 0.57 | 1.14 | 0.78 | 1.07 | 0.84 | 0.98 | 0.88 |
| P2S2 | 2.82 | 0.10 | 1.42 | 0.52 | 1.21 | 0.75 | 1.15 | 0.83 | 0.99 | 0.87 |

### 5.4. Performance Comparison of Sketching Methods within P2S-sketch

The P2S-sketch algorithm follows a sketch-and-solve [26, 82] approach and therefore the sketch size can affect the de-noising performance. In order to quantify this effect, we chose a random subject from the PPMI dataset which was acquired with a widely used 64-directions DTI protocol. A random volume (here vol. #11) from this data was first denoised with P2S and the solution **w**^∗^ obtained from it was treated as the optimal solution. Each volume of this subject contained around 960K voxels. Starting with a sketch size of 500 samples, P2S-sketch denoising was performed on the data with sketches computed using CountSketch, leverage score sampling and SRFT algorithms explained in Sec. Coresets for Regression via Matrix Sketching. The sketch size was then increased iteratively until the approximate solution 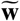 was numerically very close to **w**^∗^. The relative error for each iteration and method was computed as: 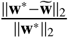.This procedure was repeated ten times to capture the variance of denoising performance since the underlying algorithms used to approximate the solution are randomized. The variance with a 95% confidence interval was plotted at each iteration (i.e., for each sketch size). We also compared the performance of the sketching methods with uniform sampling and with the deterministic choice of the rows corresponding to the top leverage scores, for the same sketching sizes. As shown in Fig. 6A, the variance of all the sampling algorithms is reduced as the sketch size increases. Uniform sampling and deterministic leverage scores perform worse than the randomized algorithms at each sketch size. While CountSketch and leverage score sampling perform approximately the same, leverage score sampling performs slightly better and offers the added advantage of interpretability. In Fig. 6C, we also empirically compare the speedup obtained from P2S-sketch in comparison with P2S. This supplements our theoretical complexity analysis in Sec. Coresets for Regression via Matrix Sketching. With experiments on three different datasets we note that the speedup obtained increases as the sketch size reduces. We also find that the speedup obtained via P2S-sketch increases in proportion to the dimensionality of the data. As one can see in Fig. 6C, the speedup on the Stanford HARDI data (shape: 81 × 106 × 76 × 160) is much more than simulated (shape: 256×256××4×63) and PPMI data (shape: 116×116×72×65). The QR decomposition computed as a part of self-supervised calibration in P2S-sketch does not add a significant computational overhead. The wall-clock time on an i7 CPU with 16GB RAM for the QR decomposition took 0.0904s on a sketch-size of 20k which amounts to 20% of the PPMI data [59] that the calibration was run on (see Fig.5). So if the experiment was to be run 10 times for calibration, the QR computation would take < 1*s*, as the subsequent runs would have fewer than 20k rows.

### 5.5. Impacts of P2S-sketch on Microstructure and Tractography

To estimate the underlying tissue **microstructure** in the living brain one typically fits a biophysical model to each voxel of the dMRI data to capture tissue heterogeneity. Diffusion kurtosis imaging (DKI) [43] is one such modeling scheme that quantifies the degree of non-Gaussian diffusion. DKI is however sensitive to noise and can often lead to fitting degeneracy in the derived maps. In Fig. 8A, we show that P2S alleviates this issue by significantly reducing the failures of model fitting in the data. We also show that P2S-sketch provides very similar estimates of the derived DKI metrics, here, radial (RK) and mean (MK) kurtosis. The same HCP 7T data was used to evaluate the DKI measures in the presence of band-like structured noise. We note that the raw noisy data shows the banded noisy artefacts on the derived DKI metrics. In order to gauge the effect of denoising, the RK and MK metrics were used to evaluate the denoising performance. Both P2S and P2S-sketch yield very similar results although P2S-sketch was trained on only 50K of the 61M samples (0.083%) of the raw data (goodness of fit comparisons in supplement). We employ the exact same procedure (with the same parameters) described in Sec. 5.2 to compare P2S-sketch against P2S for effects on tractography. In Fig. 8B we show the effect of denoising via P2S and P2S-sketch on the fiber orientation distribution (FOD) plotted via the CSD model. To de-noise via P2S-sketch, only 50K samples out of the 6M (8.3%) samples were used in the training process. Note that both P2S and P2S-sketch suppress noisy lobes uncovering the underlying fiber crossings. The generalized fractional anisotropy obtained from the constrained solid angle algorithm [1] was used as the stopping criterion of the probabilistic tracking. Next, the streamlines tracking the optic radiation and corpus callosum bundles, obtained from noisy, P2S denoised and P2S-sketch denoised data, were quantified using the fiber-to-bundle coherence metric [69] shown in Fig. 8B. The red-yellow streamlines depict the spurious and incoherent streamlines while the blue ones depict the coherent and true representative streamlines. Since the probabilistic tracking algorithm is stochastic in nature, some variability in the streamlines is expected, but both P2S and P2S-sketch yield very similar results.

**Figure 8:**
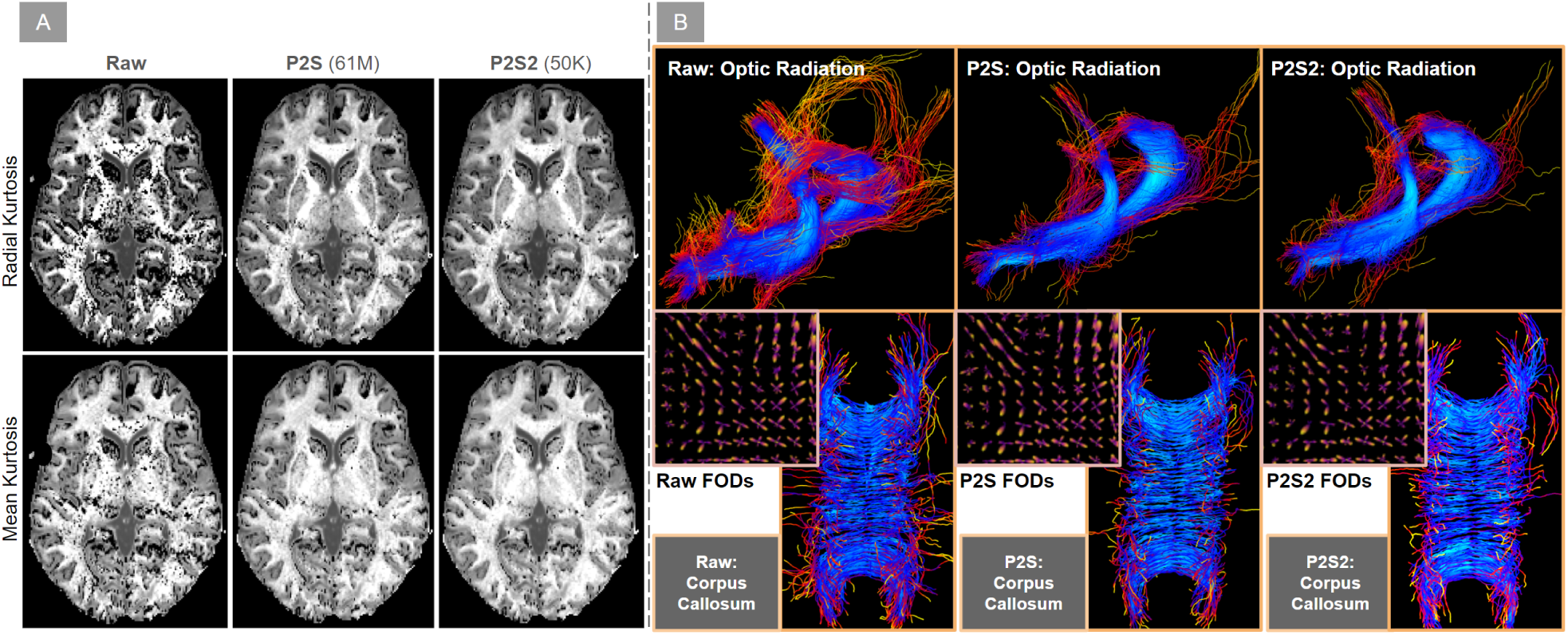
(A) Quantifies the effect of P2S and P2S-sketch denoising on HCP 7T data modeled using Diffusion Kurtosis (DKI). Derived Radial Kurtosis and Mean Kurtosis metrics are shown for both P2S and P2S-sketch. (B) We compare the performance of P2S-sketch with P2S and noisy data via fiber-to-bundle coherence quantification of the optic radiation and corpus callosum bundles. Also shows a cross-section of underlying spherical harmonic representations (FODs).

**Figure 6:**
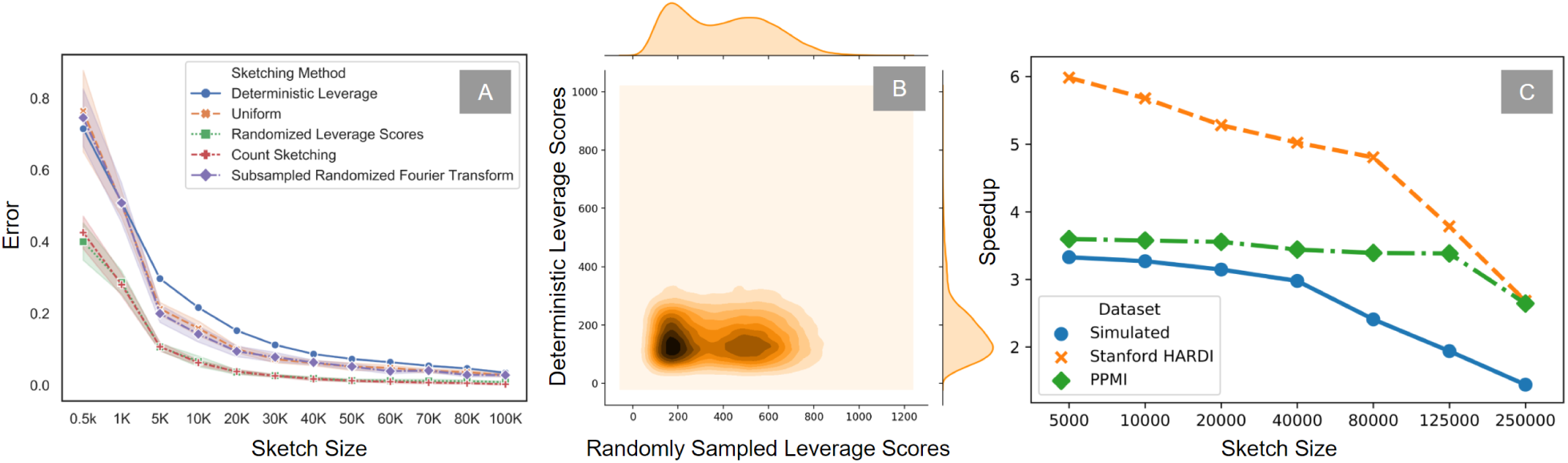
(A) We quantify the error variance in the approximation of the P2S-sketch solution vector when compared to P2S. Three different randomized algorithms (CountSketch, leverage score sampling and SRFT) are compared with uniform sampling and the deterministic choice of the rows corresponding to the largest leverage scores. (B) Joint kernel density estimate plot of the 20K voxels: randomly sampled according to leverage scores vs. deterministically chosen. (C) Shows empirical speedup comparisons (against P2S) on three different datasets, by incrementally increasing the sketch size.

### 5.6. Evaluation of P2S-sketch on Simulated Data

To compare performance on simulated data, we use a strategy similar to the one proposed in P2S (see supplement) [29, 84, 36]. The data was simulated with 2 b0 (*non-dMRI*) volumes and 60 diffusion-weighted dMRI volumes. 30 of these dMRI volumes had a b-value of 1000 *s*/*mm*^2^ and the remaining 30 with 2000 *s*/*mm*^2^. An 8-channel coil sensitivity map was used to add Gaussian noise to the real and imaginary part of each channel to simulate realistic Rician noise. Sum-of-square coil scheme was used to combine the data and the SNR was calculated in the white-matter of the b0 image. Altogether, six datasets were simulated: noise-free and SNR equal to 10, 15, 20, 25, and 30. For each SNR, we denoised the data using P2S and P2S-sketch. We compare the denoising performance of both algorithms qualitatively and quantitatively. We see that both P2S and P2S-sketch yield visually very similar results as shown in Fig. 9A. In order to quantify the difference, we compute the root mean squared error and the *R*^2^ metric between the denoised data and the ground truth at each SNR, for both P2S and P2S-sketch. The results are shown in Tab. 1. The denoising performance of P2S-sketch performance very closely approximates the denoised data obtained via P2S. Both P2S and P2S-sketch proportionally improve their performance as the SNR increases. This can also be seen in the scatter plots of signal intensities from noisy data, with P2S and P2S-sketch results shown at SNRs equal to 15 and 20 (see Fig. 9B) depicted via P2S and P2S-sketch overlap almost perfectly.

**Figure 9:**
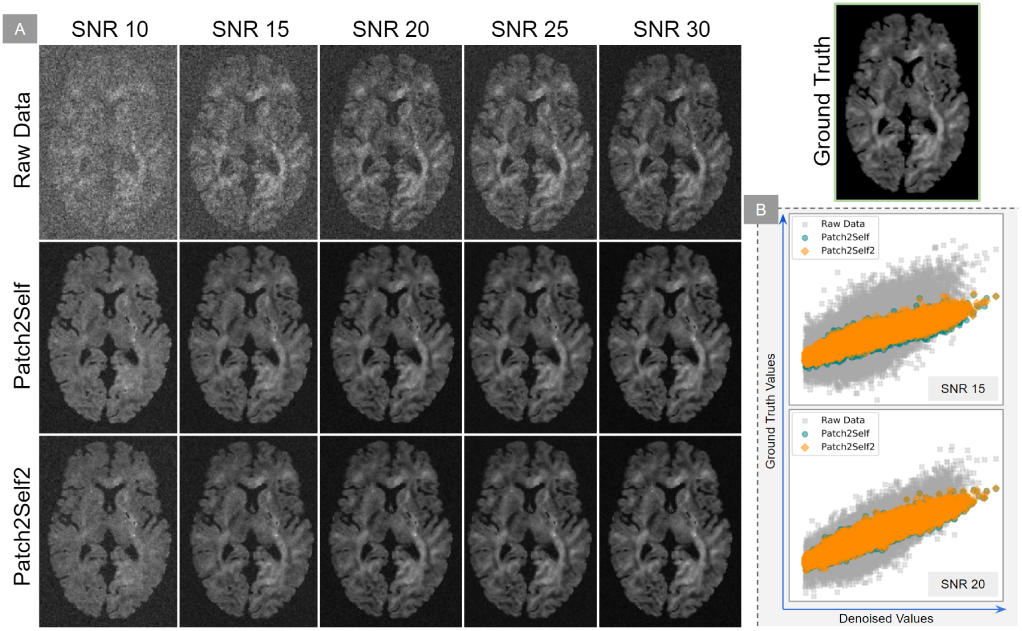
Shows (A) Qualitative comparison of denoising performance between P2S and P2S-sketch (trained on 20K samples, i.e. 7% of the data). (B) P2S-sketch closely approximates P2S via scatter plots at SNRs 15&20.

**Figure 10:**
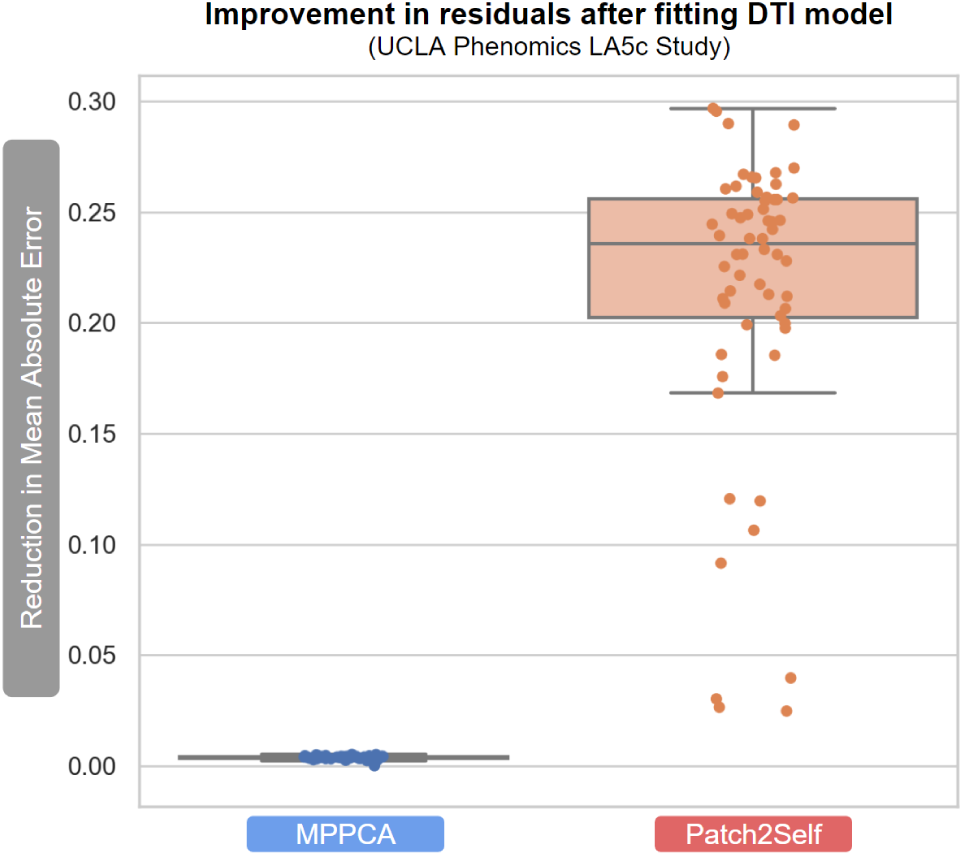
Comparison of improvement in the residuals after DTI fitting after denoising via MPPCA and P2S. Notice that P2S reduces the residual error for all 60 subjects used from the LA5c cohort in comparison with MPPCA.

### 5.7. Noise suppression and artefact removal

Visual conspicuity of the data (i.e. image quality) is crucial to any form of medical imaging, especially dMRI where the images are inherently limited by SNR. While thermal noise is known to dominate the sources of noise that corrupt the underlying signal [81], different acquisition strategies tend to induce different types of artefacts that hamper the signal structure. We show that the self-supervised setup of P2S-sketch deals with these artefacts without loss of signal corresponding to anatomical structure. In Fig. 4A, we denoised a subject **HCP 7T** dataset [80] using only 50K out of 61M (0.083%) training samples obtained via leverage score sampling. Noise mapping from this type of high-field imaging data (acquired using 7 Tesla scanner) is still under-investigated. We show that P2S-sketch suppressed band-like structured noise, which may be correlated across some volumes, but is largely uncorrelated across all 3D volumes. One of the main motives of high-field 7T scanners is to acquire data at a much higher resolution. With a zoomed-in cross-section in Fig. 4A, we show how P2S-sketch uncovers much more anatomical detail without loss of information or contrast. **Signal voids** are a common issue in MRI that occur due to certain voxels not emitting any radio-frequency signal due to a lack of activated protons in that region [83]. Since P2S, and consequently P2S-sketch, are similar to image in-painting [9], where an entire 3D volume is predicted as a combination of the rest of the volumes, this signal void can be imputed with context learned from the rest of the volumes. This setup resolves a unique issue for dMRI data which was not addressed by any other denoising algorithm in the past. In Fig. 4B we show how P2S-sketch fills the signal void present in the Stanford HARDI [72] data in gradient direction 33, without removing or smoothing the signal in the rest of the image. In Fig. 4C,D we show how P2S-sketch does not cause any signal loss even in the presence of physiological noise (porcine cardiac data) [32, 63] and ghost artefacts [70, 51], which are ubiquitous in MRI acquisitions. Note that in either case, P2S-sketch strictly only suppresses noise and does not lead to signal loss or smoothing. [In the supplement, we compare the P2S-sketch residuals with P2S].

## 6. Conclusions

This paper proposes a new method for denoising dMRI data, which is usually acquired at a low SNR, for the purpose of improving microstructure modeling, tractography, and other downstream tasks. We demonstrated that denoising by P2S outperforms the state-of-the-art random-matrix-theory-based Marchenko-Pastur method on these subsequent analyses. To enable broad adoption of this method by the MRI community, we have incorporated an efficient and unit-tested implementation of Patch2Self into the widely-used open-source library DIPY [33]. In this work we also proposed Patch2Self-sketch which performs self-supervised denoising using coresets constructed via matrix sketching, resulting in significant speedups and reduced memory usage. Our results showed that sampling-based sketching via leverage scores gave the best performance. Remarkably, leverage scores can be directly used as a statistic for interpreting influential regions hampering the denoising performance. Patch2Self-sketch will be released as a part of DIPY and a separate Pythonic matrix sketching package.

